## Supplementary materials for "Efficient rational modification of non-ribosomal peptides by adenylation domain substitution"

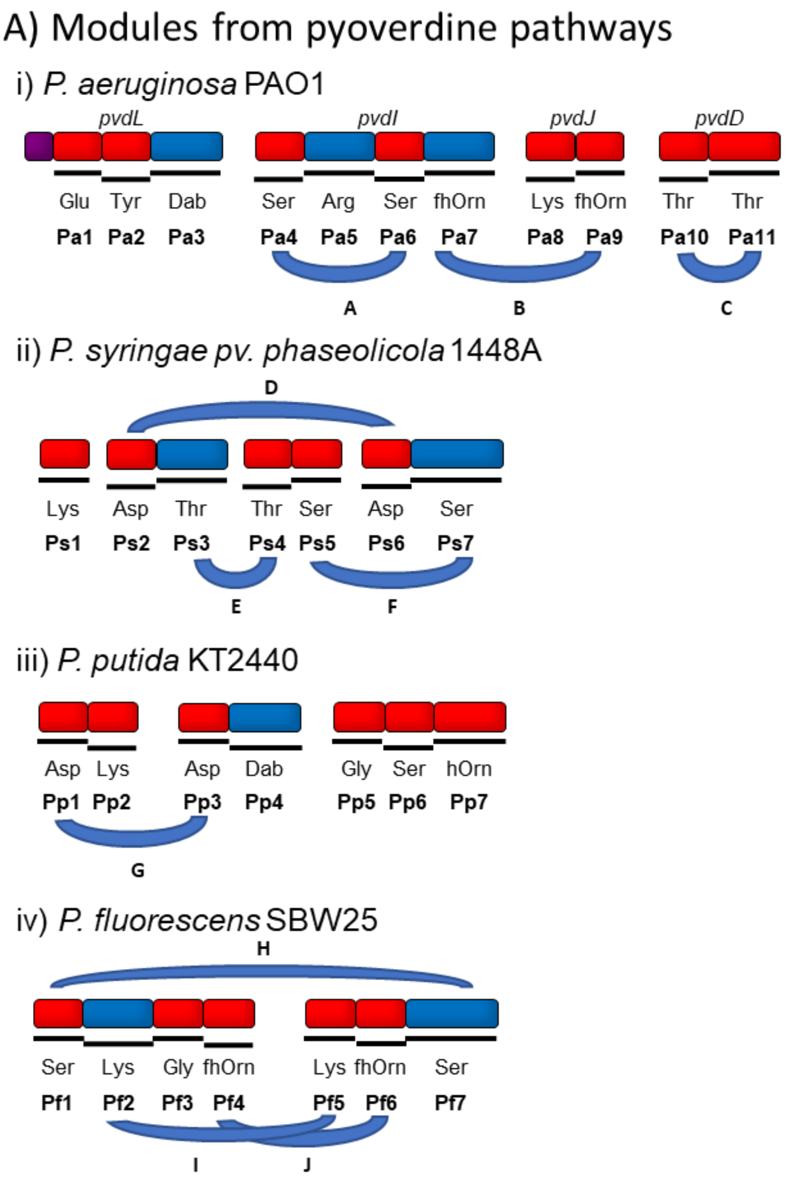


**Supplementary Figure S1:** A) NRPS modules involved in pyoverdine biosynthesis and used as a source of sequences for domain substitution and phylogenetic analysis. Modules are from NRPS pathways from i) *P. aeruginosa* PAO1, ii) *P. syringae pv. phaseolicola* 1448A, iii) *P. putida* KT2440 and iv) *P. fluorescens* SBW25. The substrate specificity and name of each module are located below the NRPS schematic. Modules within the same pathway exhibiting the same substrate specificity are linked by curved lines labelled A to J. Modules containing an $^{L}C_{L}$-domain are coloured red and those containing a $^{D}C_{L}$-domain are coloured blue.

A)

```
Pa11_Thr   QQPLALSFAQERQWFLWQLEPESAAYHIPSALRLRGRLDVDALQRSFDSLVARHETLRTR     60
Pa8_Lys    MDNLPLSYAQERQWFLWQLEPESSAYHIPTALRLRGRLDIASLQRSFAALVERHESLRTR     60
            : * **:***************:*****:*********: :*****  :**  ***:****

Pa11_Thr   FRLEGGRSYQQVQPAVSVSIER---EQFGEEGLIERIQAIVVQPFDLERGPLLRVNLLQL     117
Pa8_Lys    IARMGDEWVQVVSADVSLALEVEVQRGLDEQRLLERVEAEIARPFDLEQGPLLRVTLLEV     120
           :   *..  * *.  **:::*     . :.*: *:**::* :.:*****:******.**::

Pa11_Thr   AEDDHVLVLVQHHIVSDGWSMQVMVEELVQLYAAYSQGLDVVLPALPIQYADYALWQRSW     177
Pa8_Lys    DADEHVLVMVQHHIVSDGWSMQLMVEELVQLYAAYSQGLDVVLPALPIQYADYALWQRSW     180
             *:****:*************:**************************************

Pa11_Thr   MEAGEKERQLAYWTGLLGGEQPVLELPFDRPRPARQSHRGAQLGFELSRELVEAVRALAQ     237
Pa8_Lys    MEAGEKERQLAYWTGLLGGEQPVIELPLDHPRQPLRSYRGAQLDLELEPHLALALKQLVQ     240
           ***********************:***:*:**    :*:*****.:**. .*. *:: *.*

Pa11_Thr   REGASSFMLLLASFQALLYRYSGQADIRVGVPIANRNRVETERLIGFFVNTQVLKADLDG     297
Pa8_Lys    RKGVTMFMLLLASFQALLHRYSGQADIRVGVPIANRNRVETERLIGFFVNTQVLKADING     300
           *:*.: ************:***************************************::*

Pa11_Thr   RMGFDELLAQARQRALEAQAHQDLPFEQLVEALQPERNASHNPLFQVLFNHQSEIRSVTP     357
Pa8_Lys    RMGFDELLAQARQRALEAQAHQDLPFEQLVEALQPERSLGHNPLFQVMFNHQADSRSANQ     360
           *************************************. .*******:****:: **..

Pa11_Thr   EVQLEDLRLEGLAWDGQTAQFDLTLDIQEDENGIWASFDYATDLFDASTVERLAGHWRNL     417
Pa8_Lys    GVQLPGLSLERMEWRSSSVAFDLTLDVHEAEDGIWASFGYATDLFEASTVERLARHWQNL     420
            *** .* ** : * ..:.******::* *:*******.******:********* **:**

Pa11_Thr   LRGIVANPRQRLGEL      432
Pa8_Lys    LRGIVAEPGRPVAEL      435
           ******:* : :.**
```

Region 1

Region 2

Region 3

B)

C)

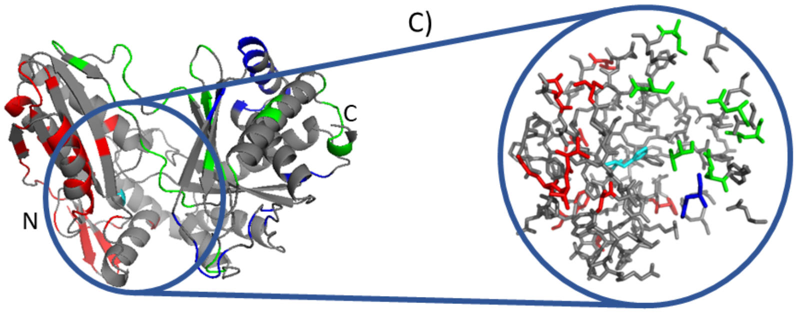


**Supplementary Figure S2:** A) Amino acid sequence alignment of the C domain from Pa11, the second module of PvdD, and the C domain from Pa8, the first module of PvdJ. The alignment is delineated into three regions defined by low homology stretches (highlighted in red, blue or green shading, respectively). Conserved motifs are underlined in bold. B) Homology model of the C domain from Pa11 identifying in colour the low-homology regions identified in panel A, and with the catalytic histidine residue shown in cyan. C). A zoomed image of the active site of the homology model.

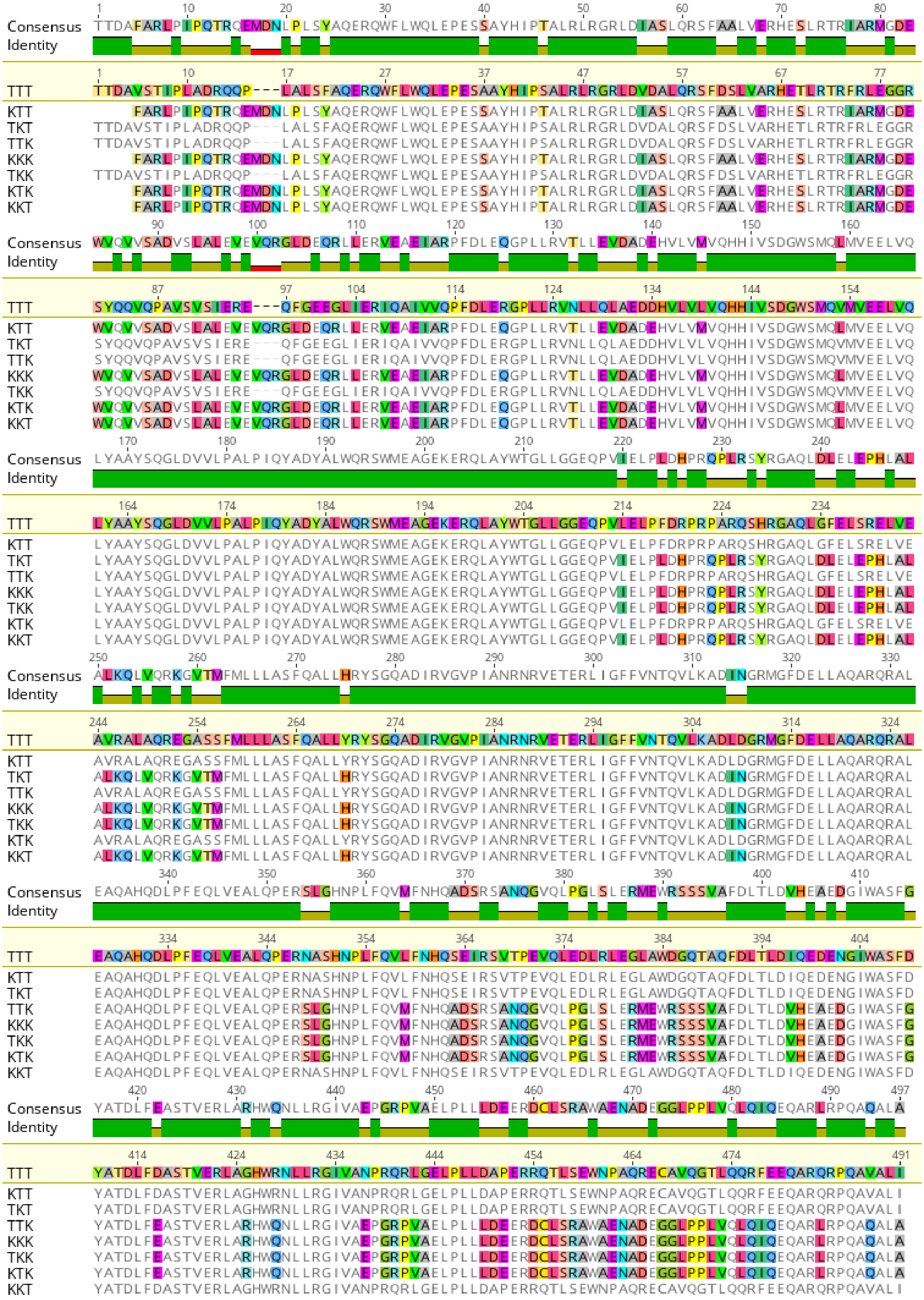


**Supplementary Figure S3:** Related to Figure 1B. Amino acid alignment highlighting the differences between shuffled C domains and the original Pa11-Thr C domain (TTT). The C domains are labelled with either a T or K for each variable region to indicate whether the region was sourced from the Pa11-Thr C domain or Pa8-Lys C domain, respectively. Image created using Geneious version 8.1 (Biomatters. Available from http://www.geneious.com).

### A) MS of shuffled C domains upstream to Thr-A domain

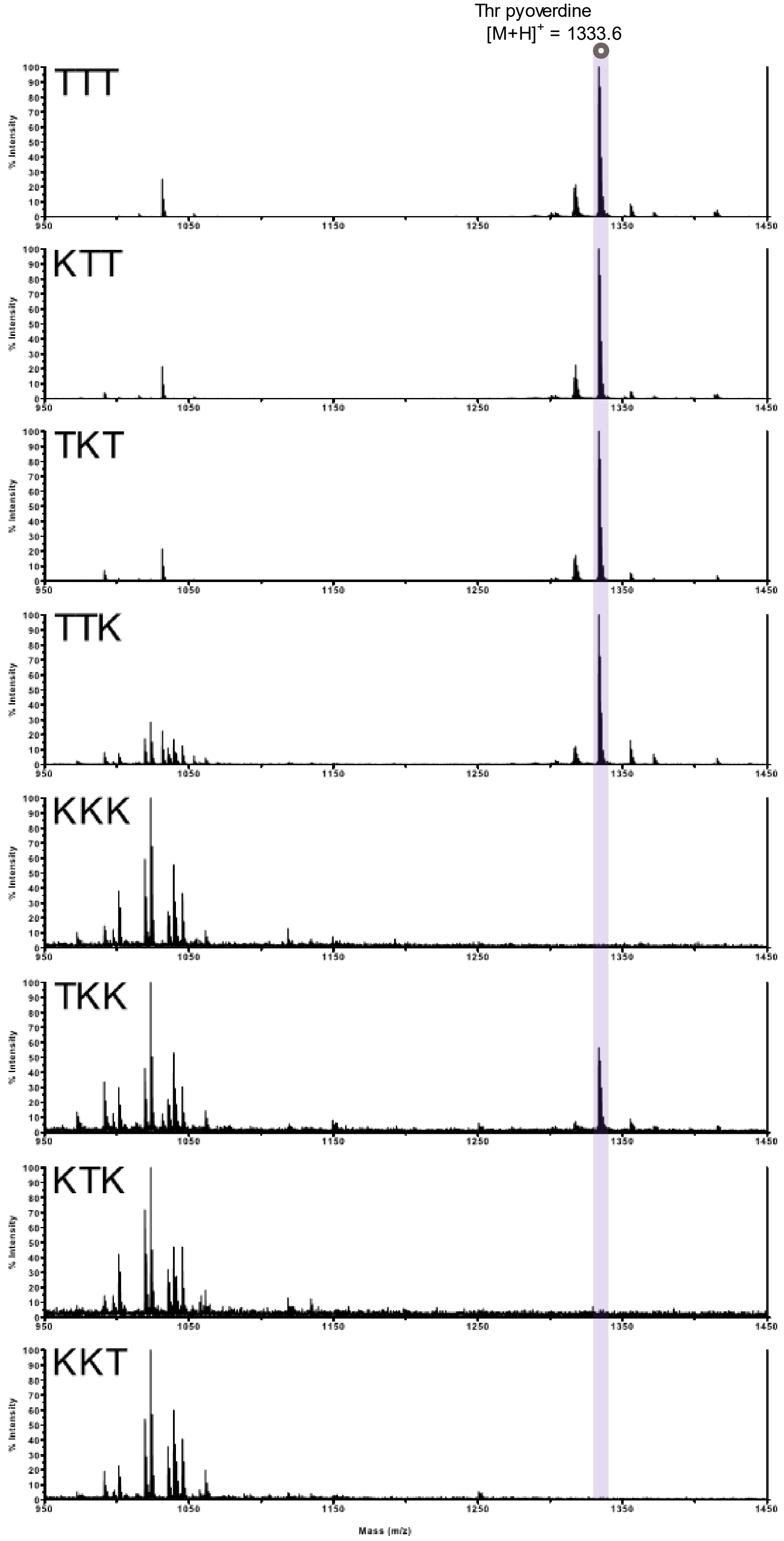

### B) MS of shuffled C domains upstream to Lys-A domain

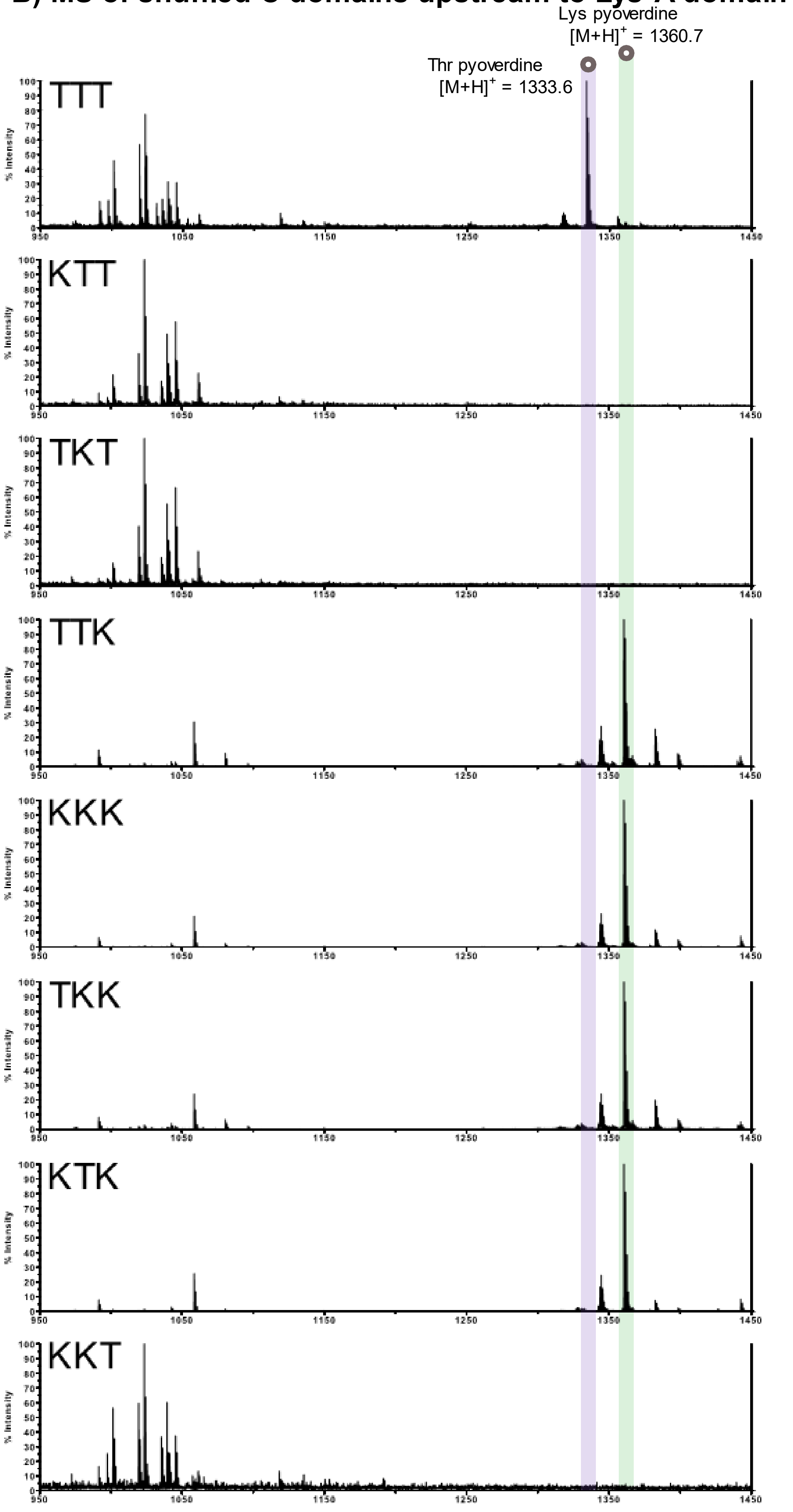

**Supplementary Figure 4.** Related to Figure 1B. Mass spectra from *P. aeruginosa* PAO1 *pvdD* strains expressing *pvdD* constructs modified in module Pa11 after shuffling the three variable regions of the Pa8 and Pa11 C domains and placing them upstream to the **(A)** Thr-specific A domain from Pa11; or **(B)** Lys-specific A domain derived from module Pa8. Spectra are labelled according to the C domains in Figure 1B of the main text. Peaks corresponding to pyoverdine with a terminal Thr (1333.6 m/z) or Lys (1360.7 m/z) are highlighted. It is noted that the C domain labelled TTT produced trace levels of WT pyoverdine, detectable only by mass spectrometry. This is consistent with previous observations for a subset of cases where a non-cognate A domain was substituted into the Pa11 module.[1] We propose that trace amounts of wild type pyoverdine reflect a non-functional Pa11 module, and stem from iterative action of the Pa10 module allowing a second L-Thr to be incorporated with low efficiency.

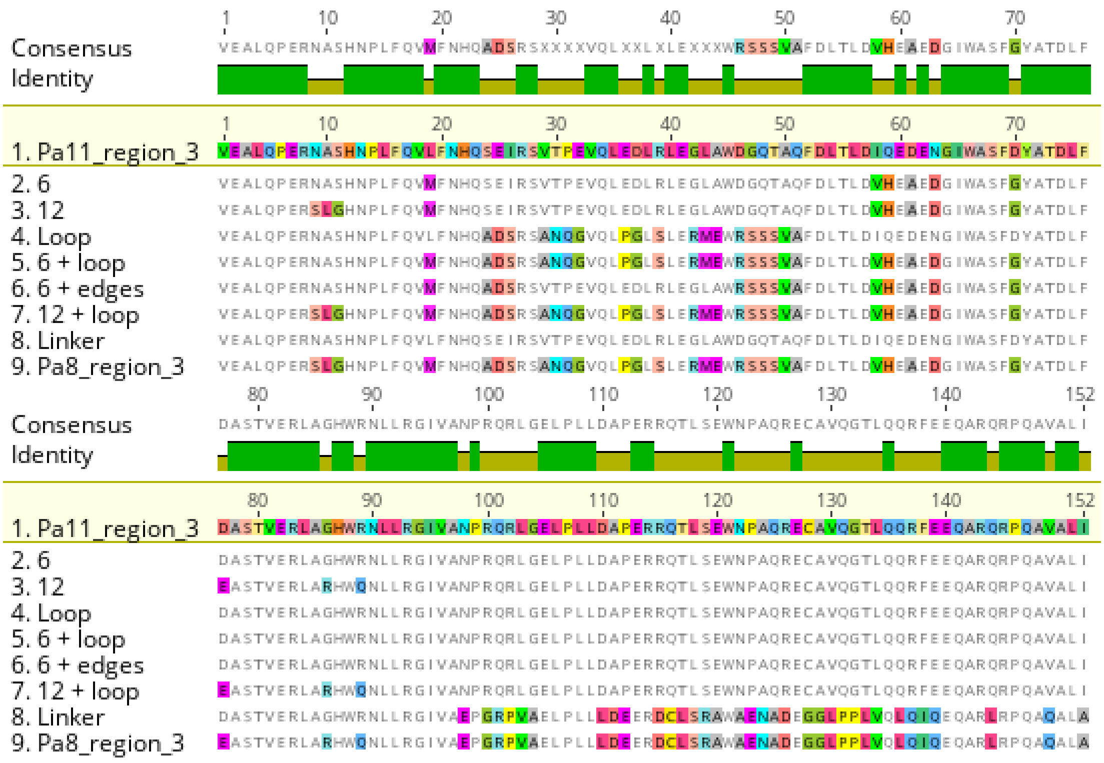

**Supplementary Figure 5:** Related to Figure 1D. Amino acid alignment highlighting the differences introduced into Pa11 during targeted mutagenesis of third variable C domain region. Alignment against the equivalent region from Pa8 is provided for reference. Image created using Geneious version 8.1 (Biomatters. Available from http://www.geneious.com).

### A) MS of region 3 mutations upstream to Thr-A domain

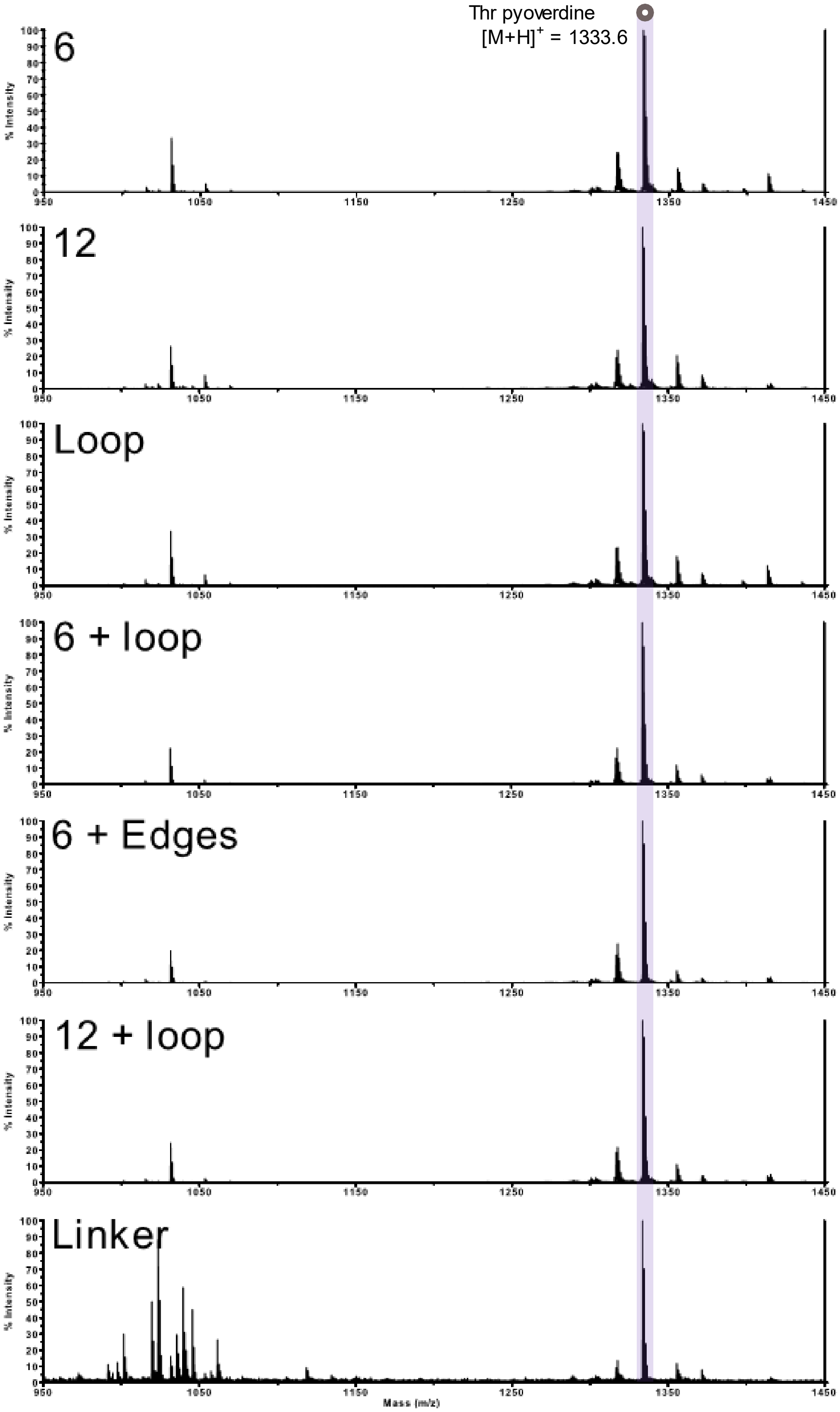

### B) MS of region 3 mutations upstream to Lys-A domain

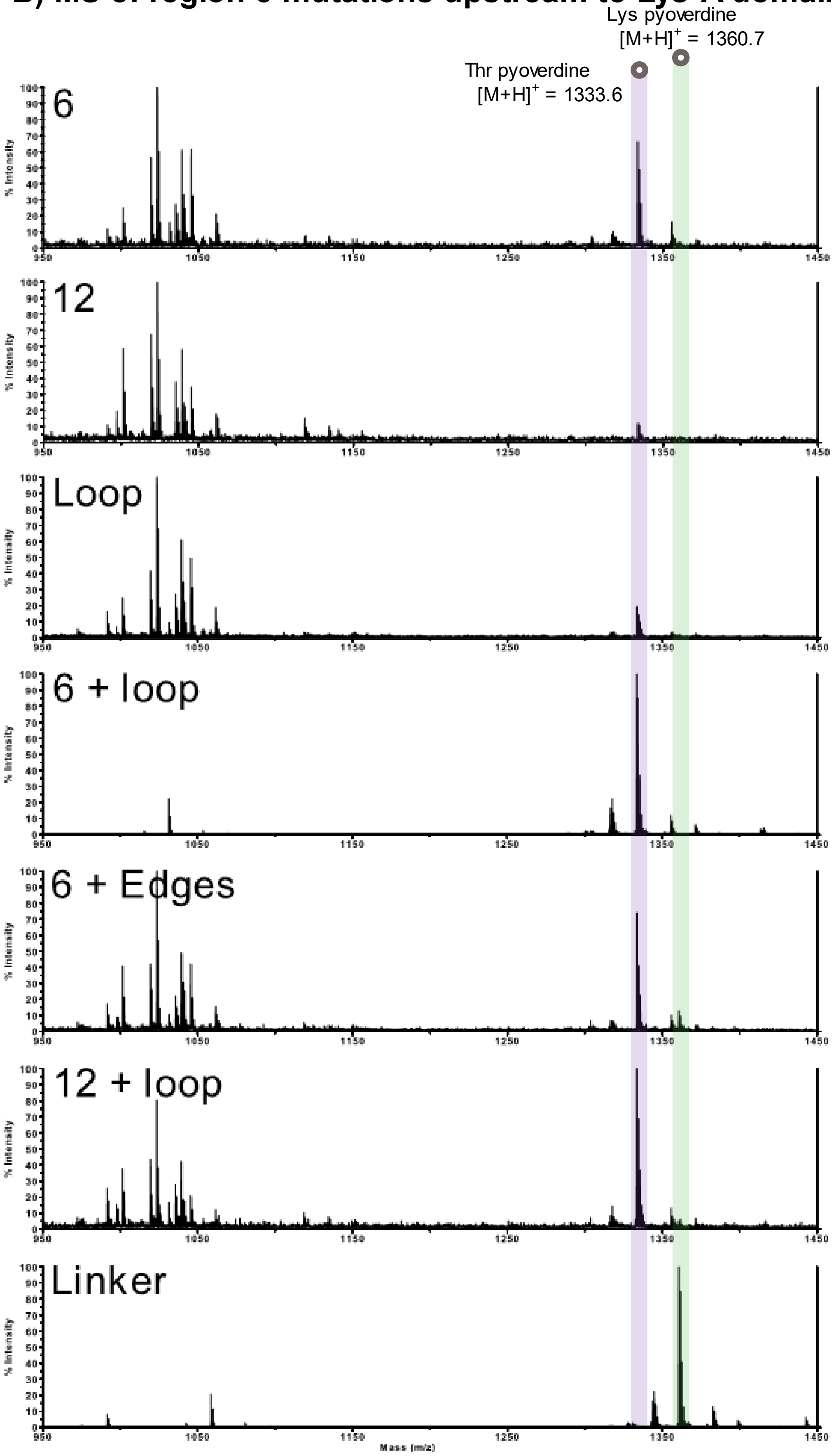

**Supplementary Figure S6.** Related to Figure 1D. Mass spectra of pyoverdine species produced by *P. aeruginosa* PAO1 strains expressing *pvdD* constructs bearing a C domain mutated in region 3 and placed upstream to the **(A)** Thr-specific A domain of Pa11, or **(B)** Lys-specific A domain from Pa8. Spectra are labelled according to the C domain order provided in Figure 1D of the main text. Peaks corresponding to pyoverdine with a terminal Thr (1333.6 m/z) or Lys (1360.7 m/z) are highlighted. Similar to Supplementary Figure S4, trace amounts of wildtype pyoverdine were detected when low-yielding C domains were upstream of the Pa8 A domain.

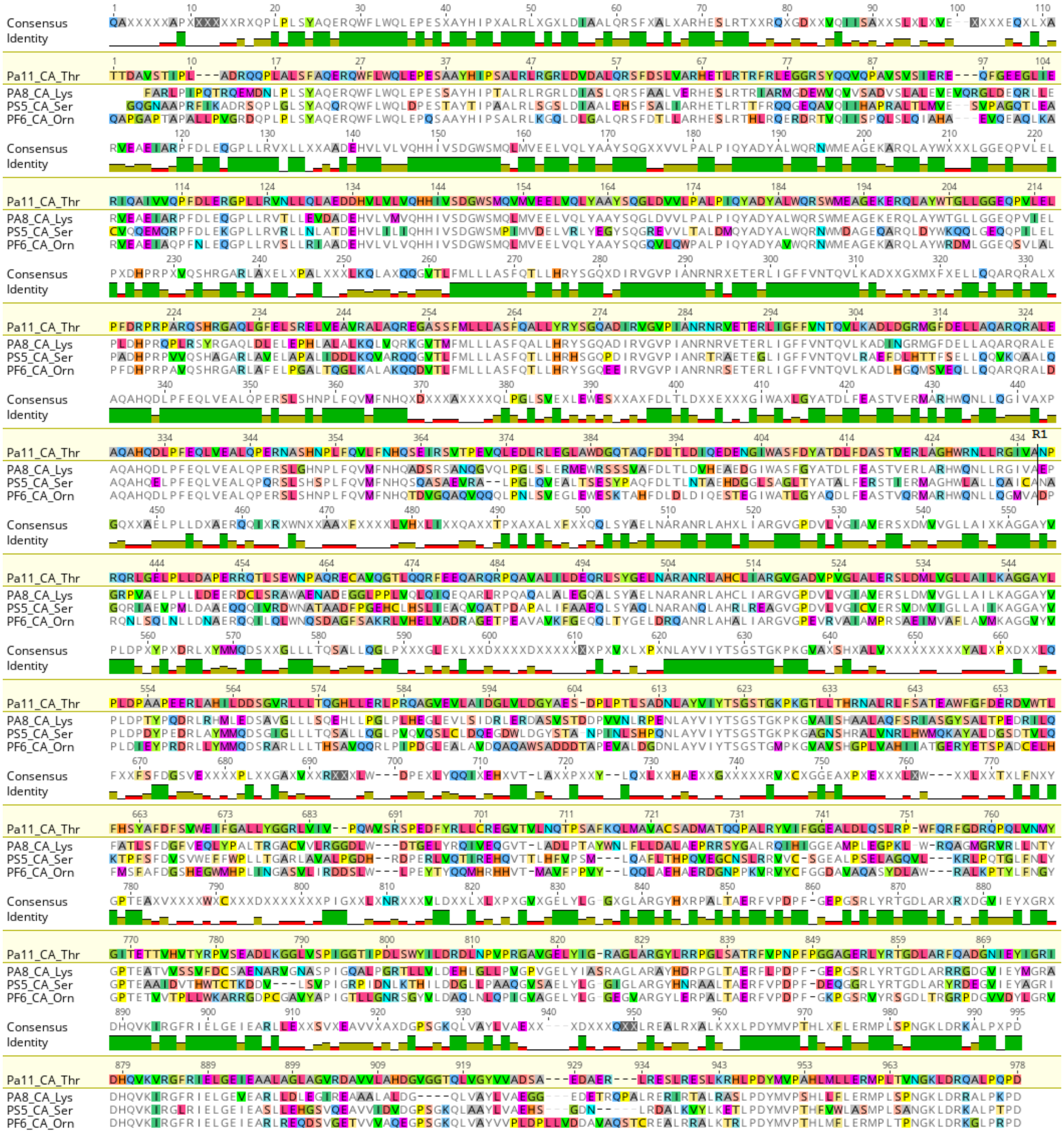

**Supplementary Figure S7.** Related to Figure 2A. Alignment of previously successful C-A domain substitutions with the C-A domains from Pa11. Each C-A domain pairing is labelled according to the module from which it was derived and its substrate specificity, as per Supplementary Figure S1. The upstream recombination point used for linker + A domain substitution is labelled R1 (A436 in Pa11_CA_Thr). Image created using Geneious version 8.1 (Biomatters. Available from http://www.geneious.com).

### MS of C-A vs linker + A domain substitutions

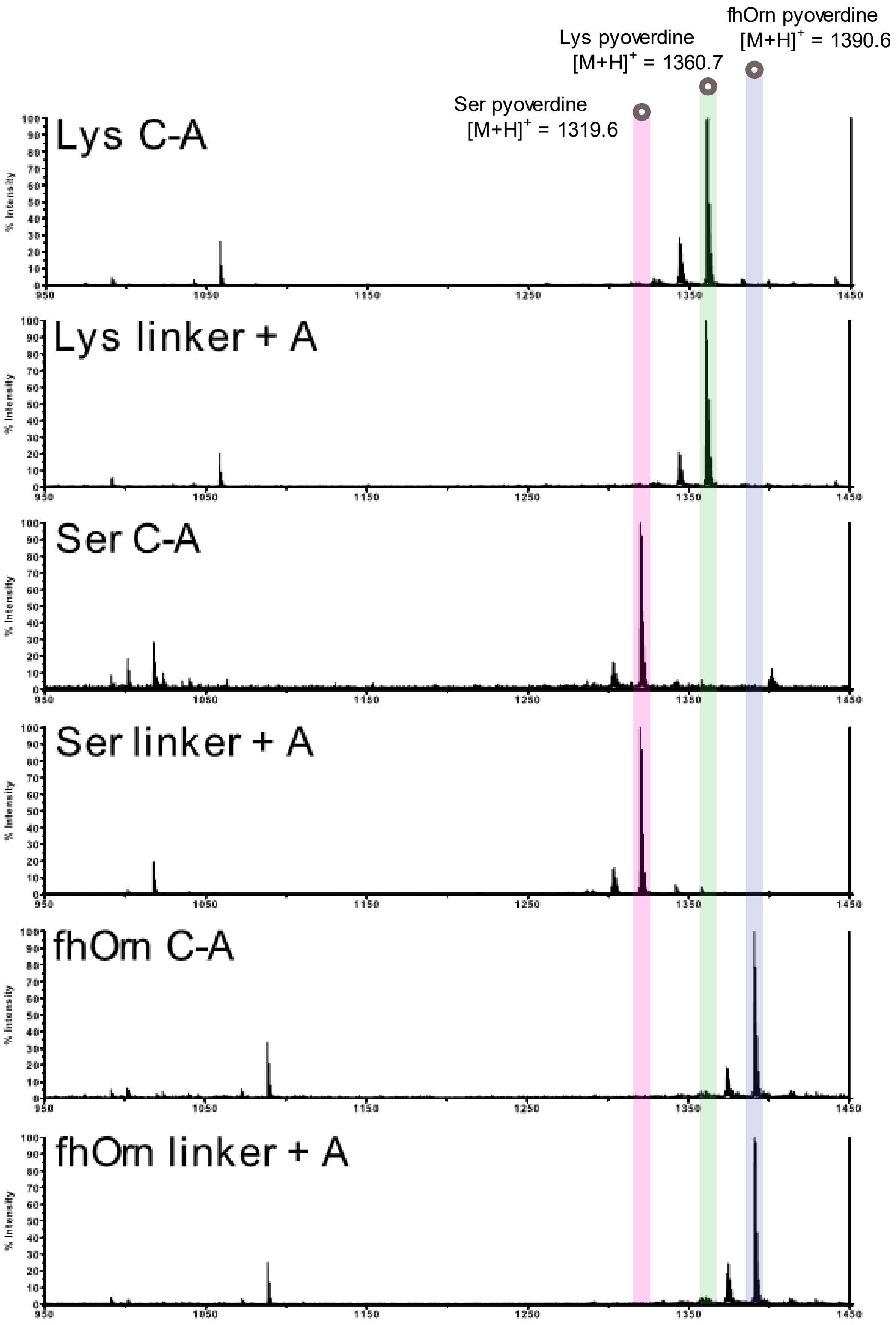


**Supplementary Figure S8.** Related to Figure 2A. Mass spectra from *P. aeruginosa* PAO1 for C-A versus linker + A domain substitutions. Peaks corresponding to pyoverdine with a terminal Lys (1360.7 m/z), Ser (1319.6 m/z) and fhOrn (1390.6 m/z) are highlighted.

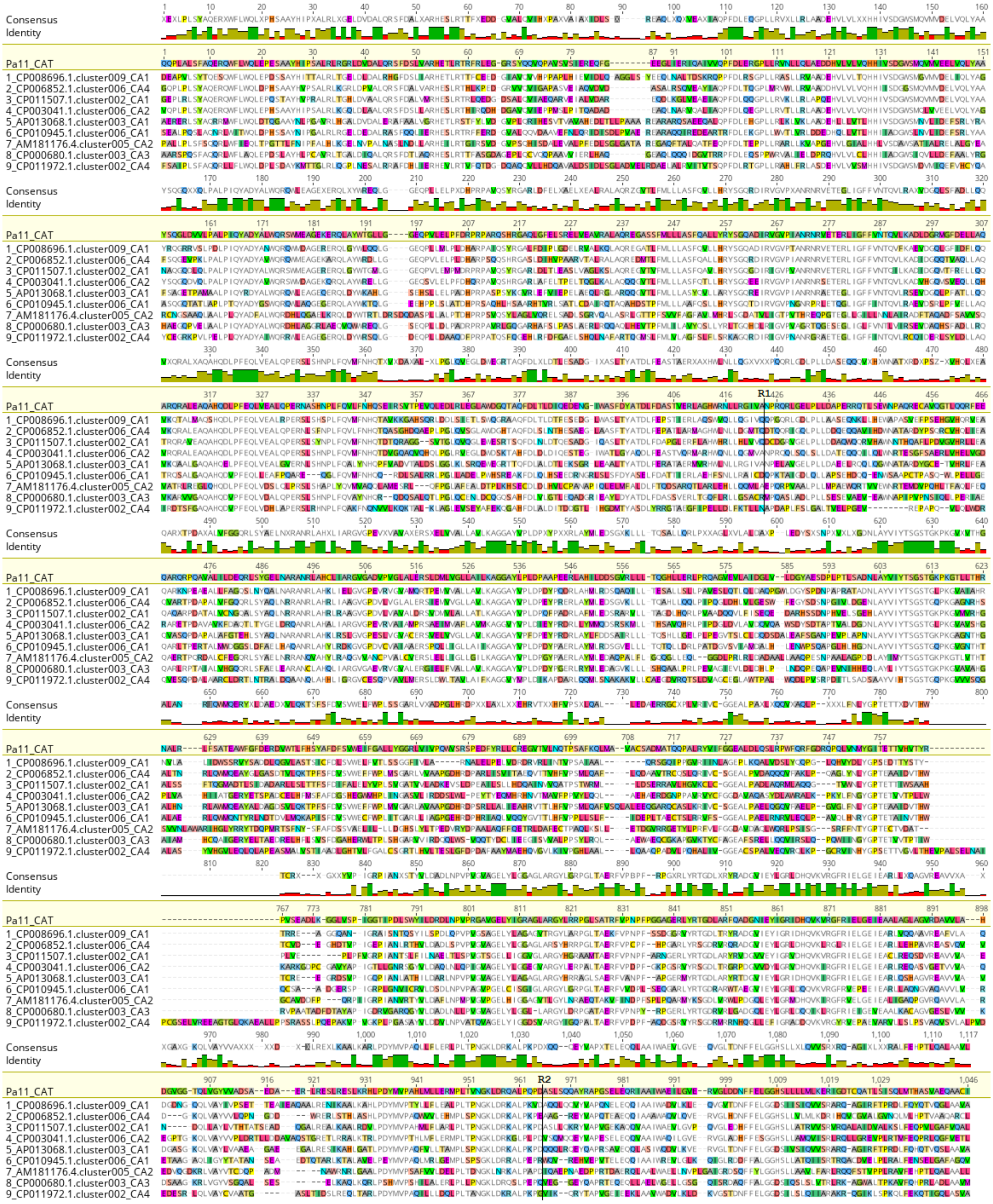

**Supplementary Figure S9.** Related to Figure 2B. Amino acid alignment highlighting differences between the C-A-T domains of module Pa11 and the C-A-T domains from modules used as a source for linker plus A domain substitution. The recombination points used for A domain substitutions are labelled R1 (N424 in Pa11_CAT) and R2 (D965 in Pa11_CAT). Image created using Geneious version 8.1 (Biomatters. Available from http://www.geneious.com).

### MS of additional linker + A domain substitutions

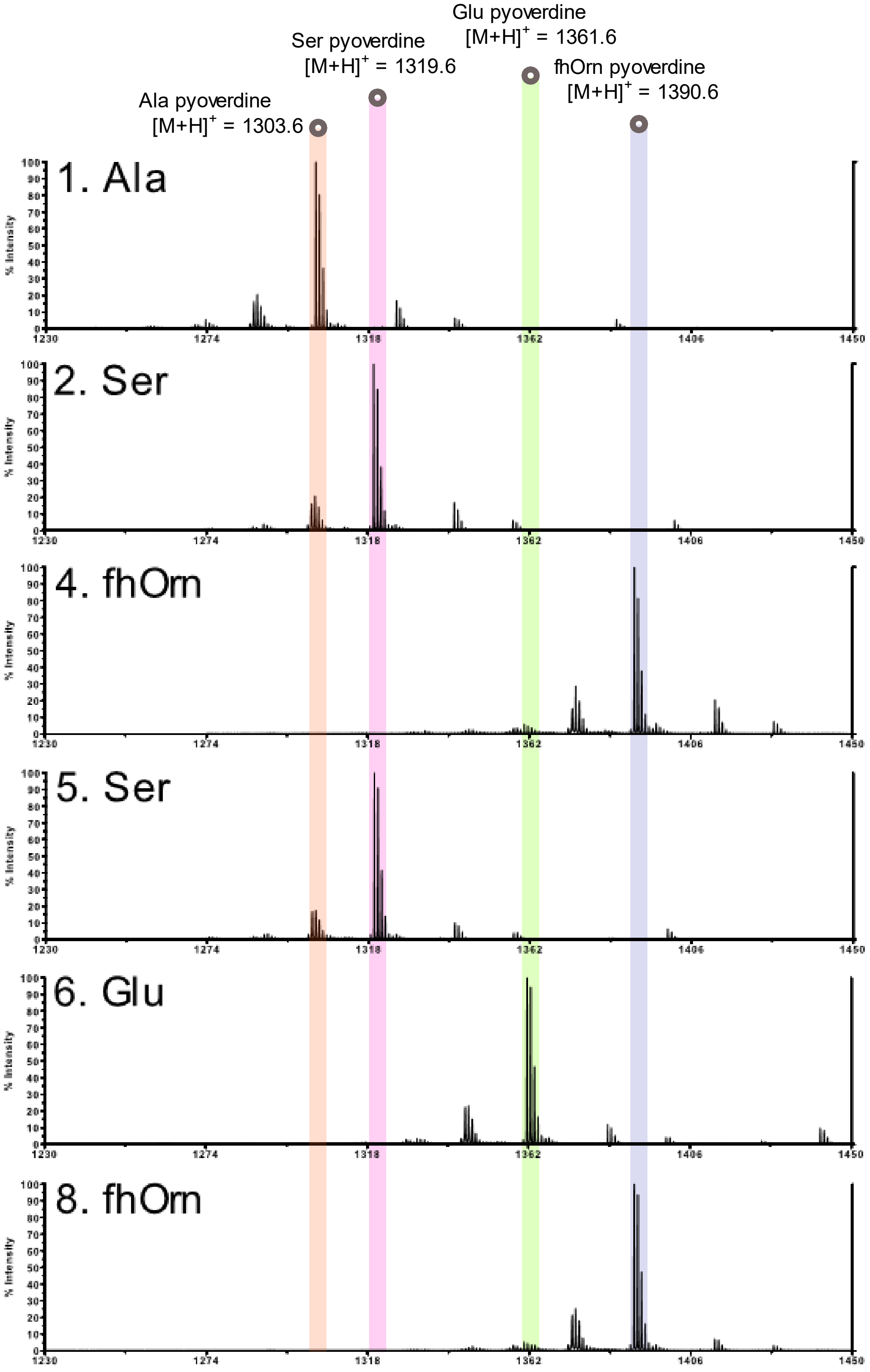


**Supplementary Figure S10.** Related to Figure 2B. Mass spectra from *P. aeruginosa* PAO1 for A domain substitution. Peaks corresponding to pyoverdine with a terminal Ala (1303.6 m/z), Ser (1319.6 m/z), fhOrn (1390.6 m/z) and Glu (1361.6 m/z).

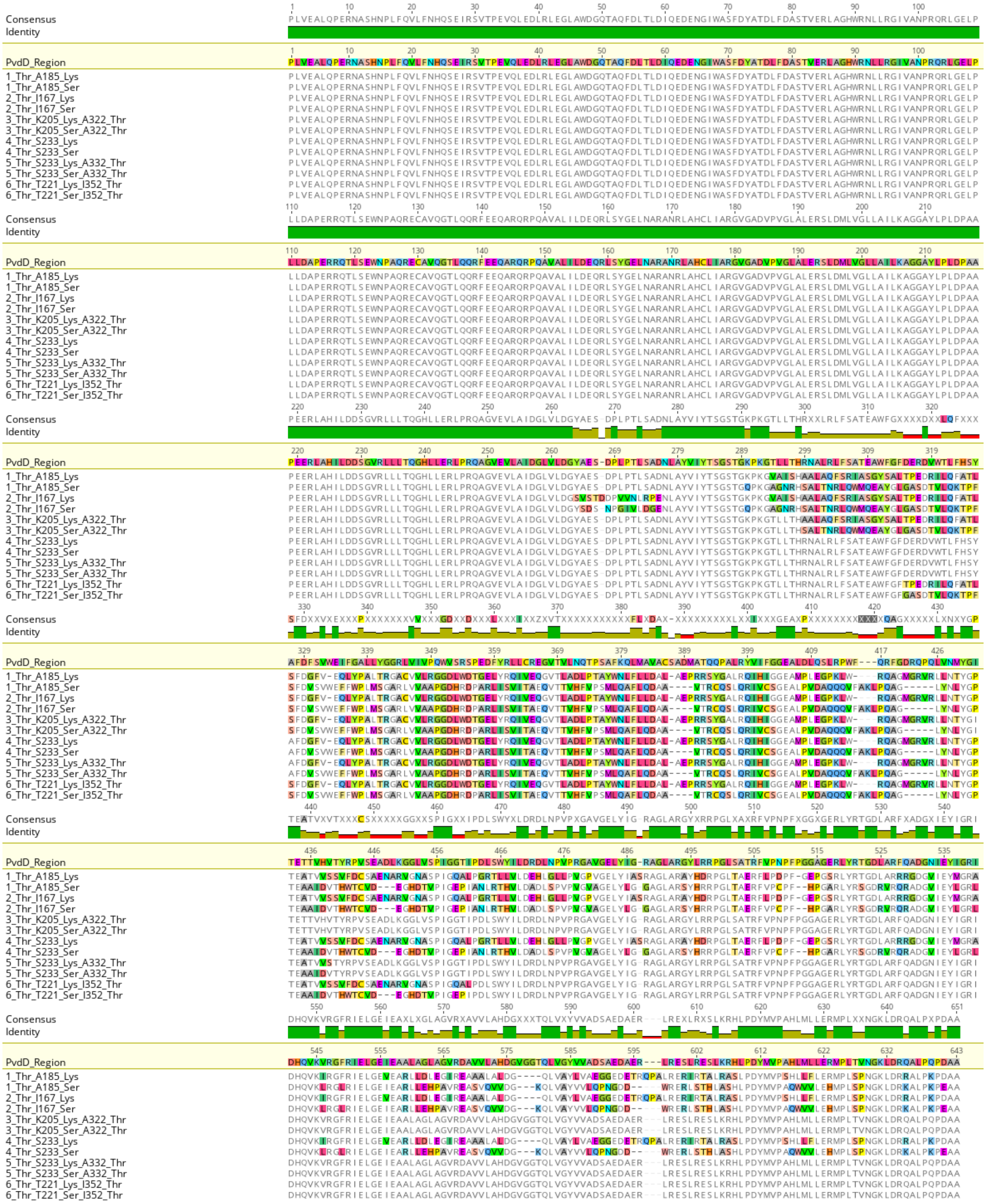


**Supplementary Figure S11.** Amino acid alignment highlighting changes to PvdD introduced by partial A domain substitutions. Each partial A domain substitution construct is labelled according to the module from which it was derived and its substrate specificity, as per Supplementary Figure S1 and Supplementary Table S1. Image created using Geneious version 8.1 (Biomatters. Available from http://www.geneious.com).

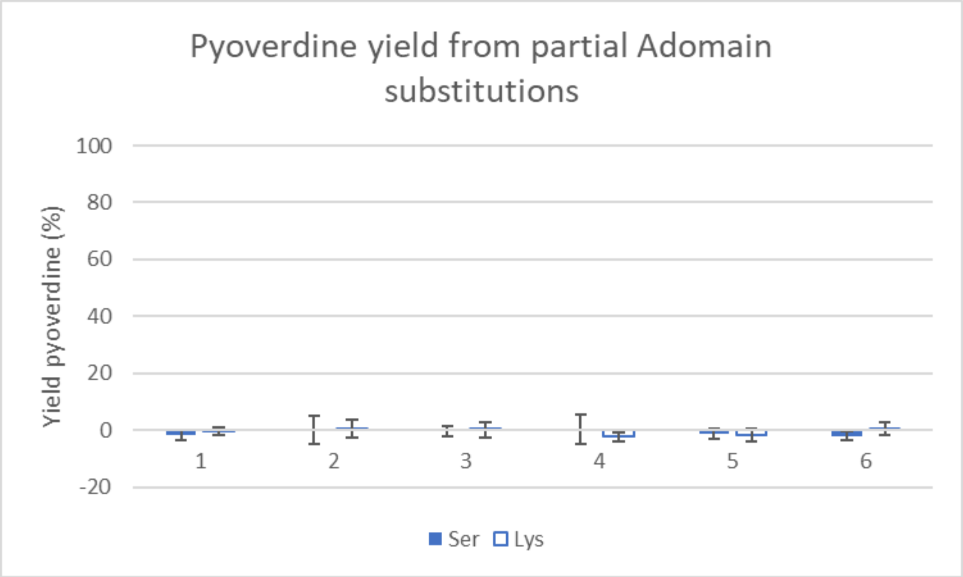


**Supplementary Figure S12.** Pyoverdine production for partial A domain substitution strains. Samples are number based on Supplementary Figure S11. Pyoverdine levels were measured by optical density at 400 nm, and error bars represent the standard deviation from six independent replicates.

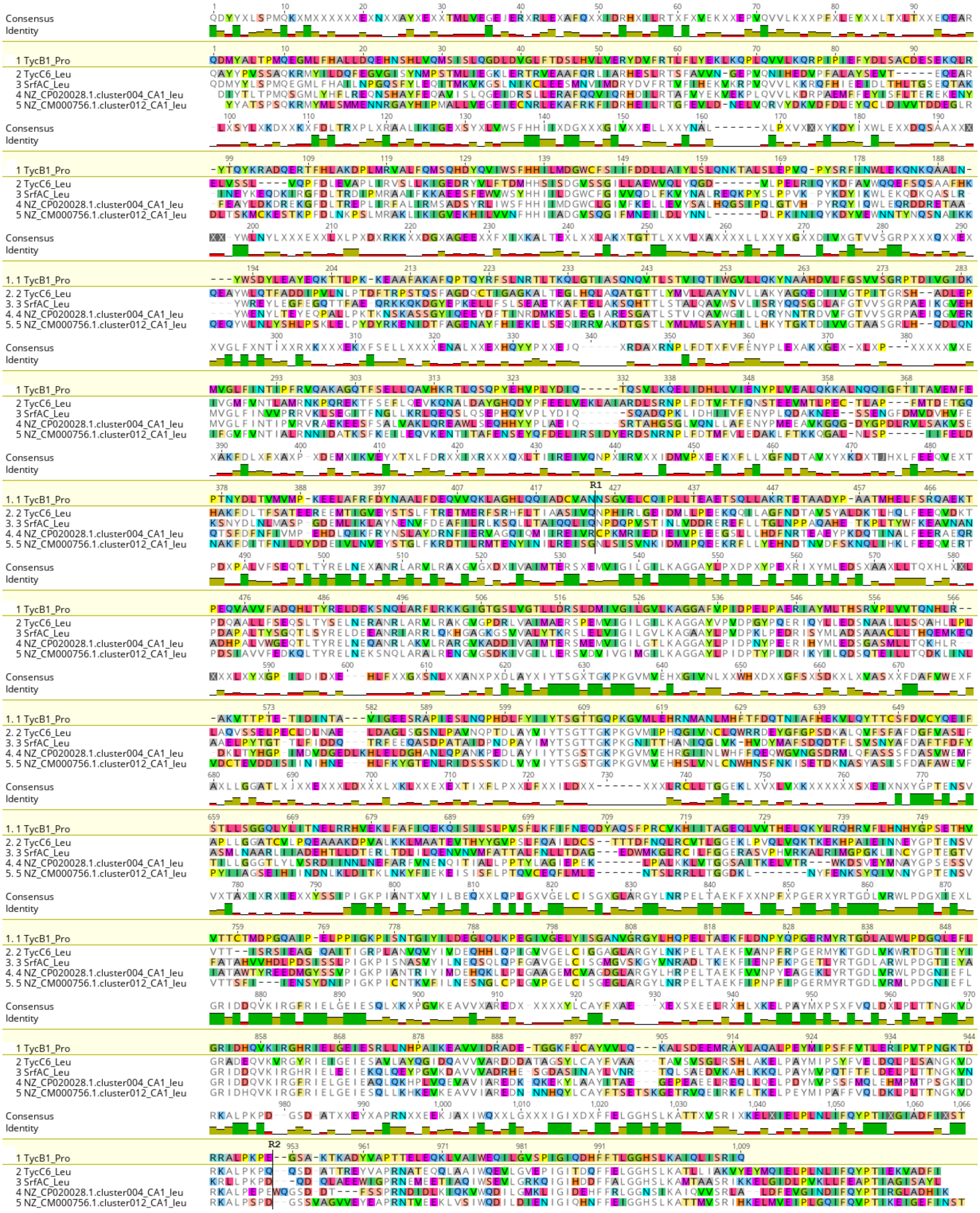


**Supplementary Figure S13:** Related to Figure 5. Amino acid alignment highlighting the differences between the C-A-T domains of ProCAT and the C-A-T domains from the Leu-specifying modules used for linker + A domain substitutions. The recombination points used for A domain substitutions are labelled R1 (N424 in TycB1_Pro) and R2 (E952 in TycB1_Pro). Image created using Geneious version 8.1 (Biomatters. Available from http://www.geneious.com).

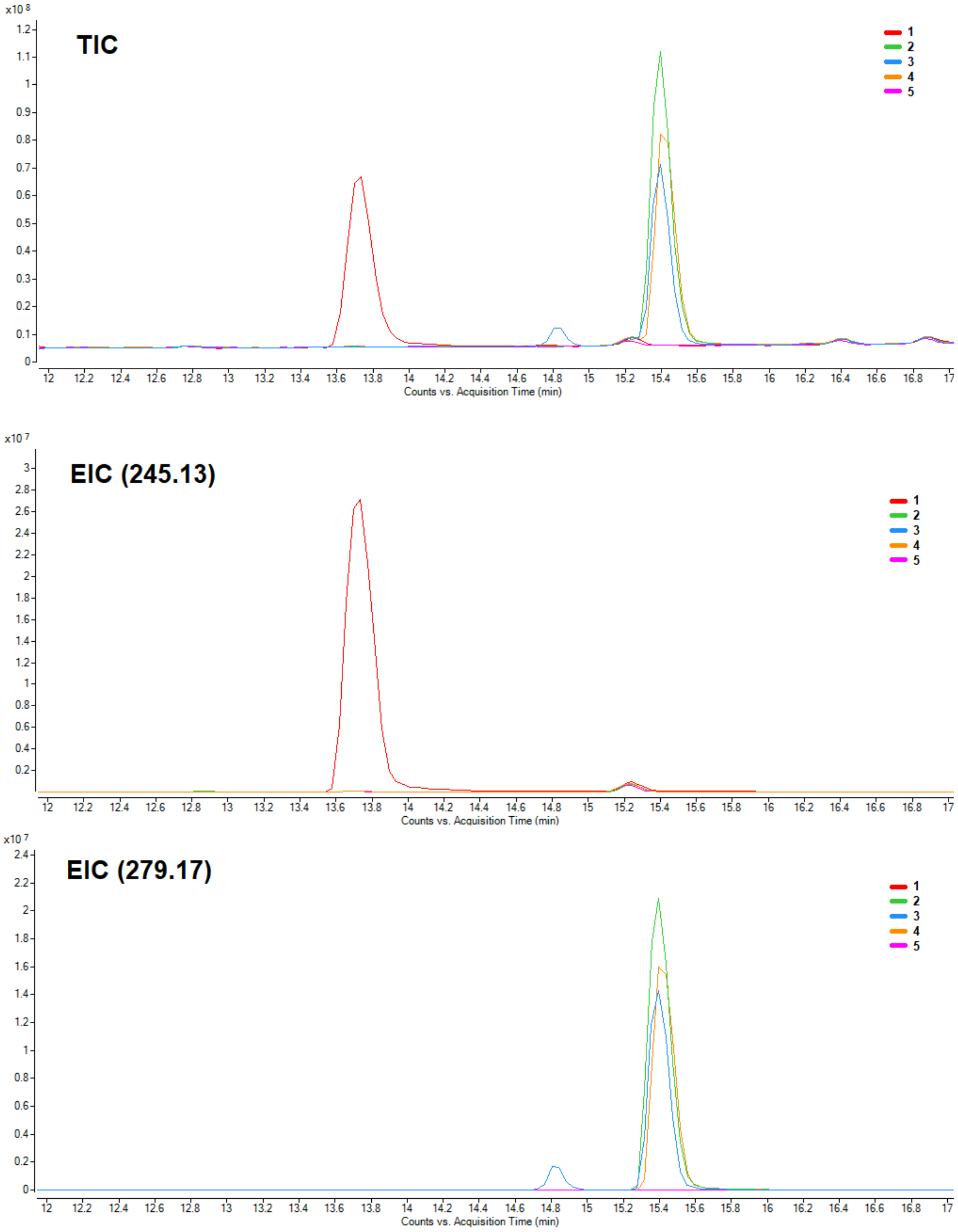


**Supplementary Figure S14.** Related to Figure 5. Mass spectrometry traces for A domain substitution strains using the PheATE/ProCATTe system. Total ion count and the EICs for Phe-Pro DKP ($[M+H]^+$ = 245.13) and Phe-Leu ($[M+H]^+$ = 279.17) are shown. Traces are labelled 1 to 5 as per Figure 5.

**Supplementary Table S1** - Substrate specificity predictions for each A domain substituted into PvdD, together with the name of each cluster, and amino acid identity shared between the C and A domains of each module and the corresponding domains from module Pa11-Thr.

| | Specificity predictions from the AntiSMASH database | | | | Cluster name* | Amino acid identity to Pa11-Thr (%)** | |
|---|---|---|---|---|---|---|---|
| | NRPS Predictor2 SVM | Stachelhaus code | Minowa | Consensus | | C domain | A domain |
| 1 | ala | ala | ala | ala | CP008696.1.cluster009_CA1 | 66.57 | 51.32 |
| 2 | ser | ser | ser | ser | CP006852.1.cluster006_CA4 | 72.24 | 50.54 |
| 3 | gly | gly | gly | gly | CP011507.1.cluster002_CA1 | 71.94 | 47.58 |
| 4 | hydrophilic | orn | orn | orn | CP003041.1.cluster006_CA2 | 73.43 | 47.01 |
| 5 | ser | ser | ser | ser | AP013068.1.cluster003_CA1 | 58.51 | 50.64 |
| 6 | glu | glu | ser | glu | CP010945.1.cluster006_CA1 | 53.61 | 48.39 |
| 7 | asp,asn,glu,gln,aad | asp | orn | nrp | AM181176.4.cluster005_CA2 | 42.99 | 43.98 |
| 8 | hydrophilic | trp | orn | nrp | CP000680.1.cluster003_CA3 | 53.13 | 47.39 |
| 9 | asp | asn | asp | asp | CP011972.1.cluster002_CA4 | 53.73 | 39.48 |

*Domains were named according to the cluster name from the AntiSMASH database,[2] and a number based on the order the CA-domains appeared in the GBK file.

**C domains were trimmed to the C1 and C7 motifs, and A domains were trimmed to the A1 and A10 motifs inclusive. Domains were aligned using MUSCLE,[3] and the resulting alignment used to calculate percent identity to the corresponding sequence from module Pa11.

**Supplementary Table S2** - Substrate specificity predictions for each A domain substituted into ProCATTe, together with the name of each cluster, and amino acid identity shared between the C and A domains of each module and the corresponding domains from ProCATTe.

| | Specificity predictions using Stachelhaus code | Cluster name* | Amino acid identity to Pa11-Thr (%)** | |
|---|---|---|---|---|
| | | | C domain | A domain |
| 2 | Leu | TycC6_Leu | 22.98 | 43.11 |
| 3 | Leu | SrfAC_Leu | 44.08 | 40.35 |
| 4 | Leu | NZ_CP020028.1.cluster004_Leu_CA1 | 43.07 | 47.02 |
| 5 | Leu | NZ_CM000756.1.cluster012_Leu_CA1 | 21.82 | 47.64 |

*Domains were named according to the cluster name from the AntiSMASH database.[2]

**C-domains were trimmed to the C1 and C7 motifs, and A domains were trimmed to the A1 and A10 motifs inclusive. Domains were aligned using MUSCLE,[3] and the resulting alignment used to calculate percent identity to the corresponding sequence from ProCAT.
